# SurvMarker: An R Package for Identifying Survival-Associated Molecular Features Using PCA-Based Weighted Scores

**DOI:** 10.64898/2025.12.31.697184

**Authors:** Dona Hasini Gammune, Tongjun Gu

## Abstract

**Summary:** SurvMarker is an R package for identifying survival-associated features in high-dimensional molecular data using PCA-based weighted scoring. The method aggregates feature loadings across survival-associated principal components (PCs) and evaluates feature significance against a feature-specific empirical null distribution, enabling stable, parsimonious, and statistically calibrated prognostic feature selection.

**Availability and Implementation:** SurvMarker is implemented in R and distributed under the MIT License (MIT). The package includes comprehensive documentation, a reference manual, and a reproducible workflow example, and is provided as Supplementary Material. Source code and documentation are openly available at the GitHub repository https://github.com/tjgu/SurvMarker.

## Introduction

High-dimensional transcriptomic RNA profiling has become central to biomarker discovery and prognostic modeling in cancer and other complex diseases. In survival analysis settings, dimensionality reduction is often required to stabilize inference, reduce multiple-testing burden, and retain biologically meaningful signal. Principal component analysis (PCA) is widely used for this purpose, as it captures dominant patterns of variation in expression data and provides orthogonal latent representations suitable for downstream survival modeling.

Despite its widespread use, there is no principled and widely accepted approach for selecting molecular features from principal components (PCs). The most common strategy is to select a fixed number of top-loading features from each PC (Pollen et al., 2014), an approach also adopted in our earlier work (Gammune et al., 2025). While effective, this strategy has several limitations. First, it relies on an arbitrary per-PC cutoff. Second, PCs are treated independently, without integrating information across multiple survival-associated PCs. Consequently, features with moderate but consistent contributions across several PCs, potentially reflecting biologically meaningful and prognostic relevant signals, may be overlooked.

To address these limitations, we developed SurvMarker, an R package that implements a novel, data-driven PCA-based feature selection framework for survival analysis. Rather than selecting a fixed number of features from individual PCs, SurvMarker aggregates loading information across all survival-associated PCs and explicitly anchors feature importance to survival relevance.

Within this framework, PCA is performed on a normalized high-dimensional expression matrix, and survival-associated PCs are identified using Cox proportional hazards models, with optional adjustment for clinical covariates. Feature importance is quantified by aggregating loadings across all survival-associated PCs, weighted by the variance explained by each PC, yielding a single interpretable score for each gene that reflects its overall contribution to survival-relevant latent structure. Statistical significance is assessed using an empirical null distribution constructed from non–survival-associated PCs, enabling estimation of empirical *p*-values and false discovery rates.

To our knowledge, this loading-based feature scoring strategy has not been reported previously. The most closely related methodology is supervised PCA (Bair et al., 2006), which links dimensionality reduction to outcome-guided component selection. However, supervised PCA typically derives feature importance from a single supervised component or from direct correlations between features and outcome-guided latent variables. In contrast, our approach aggregates information across multiple survival-associated PCs, incorporates variance-based weighting, and uses an empirical null framework for inference. This results in a more stable, interpretable, and statistically grounded ranking of prognostic features, particularly in settings where survival signal is distributed across multiple latent dimensions.

The proposed framework is implemented in the SurvMarker R package, provides end-to-end functionality for PCA-based survival screening, feature scoring, empirical null construction, and visualization. While motivated by transcriptomic expression data, SurvMarker is applicable to any normalized high-dimensional molecular matrix, including miRNA expression, proteomic and metabolic measurements, and DNA methylation ratios.

The remainder of this paper describes the SurvMarker framework and its implementation. Figure 1 presents representative outputs generated using real-world biological data, and additional methodological details and extended analyses are provided in the Supplementary Material.

**Fig 1.**
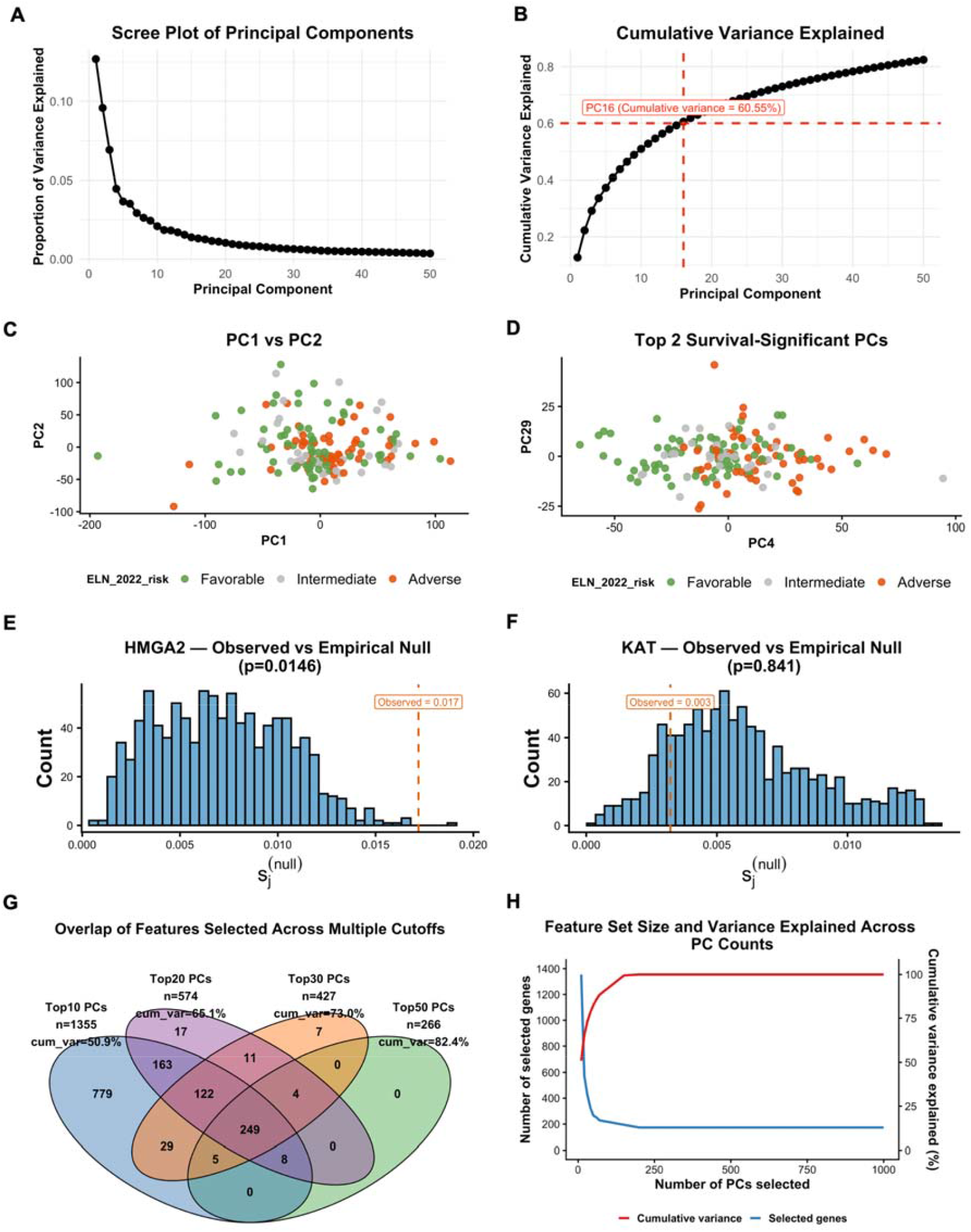
Survival-associated PCA-based feature scoring and visualization framework implemented in SurvMarker. (A) Scree plot showing the proportion of variance explained by each principal component (PC). (B) Cumulative variance explained across PCs. (C) Scatter plot of PC1 versus PC2. (D) Scatter plot of the top two survival-associated PCs. (E–F) Feature-specific empirical null distributions of the weighted score for a prognostically significant gene (HMGA2) and a non-significant gene (KAT), respectively. (G) Overlap of selected features across multiple PC thresholds. n denotes the number of features selected at different thresholds. (H) Trade-off between feature set size and cumulative variance explained as the number of

### SurvMarker: Implementation and Results

SurvMarker is implemented as an R package and distributed under the MIT License (MIT). R was chosen due to its widespread adoption in the bioinformatics community, rich ecosystem of statistical and visualization tools, and seamless integration with existing transcriptomic and survival analysis workflows. The package is designed to support reproducible analyses and straightforward incorporation into established genomic pipelines.

PCA-based feature selection in high-dimensional survival analysis involves several nontrivial methodological choices, including the number of PCs to evaluate, how information should be aggregated across PCs, and how statistical significance should be assessed while controlling false discoveries. SurvMarker provides a transparent and flexible framework that exposes these decisions through well-documented parameters, enabling users to tune analyses while maintaining robust default behavior. Representative outputs are shown in Figure 1, while the complete feature scoring methodology is illustrated in Supplementary Figure 1.

### Core workflow and feature scoring

A typical SurvMarker analysis is performed using a single function call:

~~~
pca_based_weighted_score(
X = X, time = time, status = status, covar = covariates,
n_pcs = 50, pc_fdr_cutoff = 0.05, feature_fdr_cutoff = 0.05,
null_B = 500, scale_pca = TRUE, use_abs_loadings = TRUE,
store_null = TRUE, seed = 1)
~~~

The input matrix, `X`, is a normalized high-dimensional dataset with samples in rows and features (e.g. genes or miRNAs) in columns. Survival follow-up time and event status are provided via `time` and `status` (1 = event, 0 = censored), respectively. Optional clinical covariates (`covar`) may be included to adjust Cox proportional hazards models for PC–survival associations (e.g., age, sex, and European LeukemiaNet (ELN) 2022 risk).

SurvMarker provides two complementary strategies for determining the number of PCs evaluated. Users may specify a fixed number via `n_pcs`, or alternatively select the minimum number of PCs (`n_pcs`) required to explain a desired fraction of variance via `cumvar_threshold`, with `max_pcs` (default 50PCs) acting as a safety cap. Survival association is tested for each PC, and multiple testing is controlled using False Discovery Rate (FDR) across PCs (`pc_fdr_cutoff`, default 0.05), defining the set of survival-associated components.

Rather than selecting a fixed number of top-loading features per PC, SurvMarker aggregates feature loadings across all survival-associated PCs to compute a single weighted score *S*_*j*_ for each feature. Loadings can be taken in absolute value (`use_abs_loadings = TRUE`) and are weighted by the variance explained by each PC, prioritizing features that contribute consistently to survival-relevant latent structure. Statistical significance is assessed using an empirical null distribution constructed by repeatedly sampling from non–survival-associated PCs (`null_B`, default 500). Empirical *p*-values and Benjamini–Hochberg (BH)-adjusted *p*-values are computed, and prioritized features are selected using a user-defined FDR threshold (`feature`_`fdr_cutoff`, default 0.05).

The returned object includes a ranked feature table (`feature_table`) containing each feature’s loadings on survival-associated PCs, the aggregated score *S*_*j*_, empirical *p*-value, and BH-adjusted *p*-value, as well as PC-level summaries of variance explained and survival associations (`pc_table`). For reproducibility and diagnostic purposes, SurvMarker optionally stores the full null score matrix (`null_scores` when `store_null = TRUE`), PC scores, and PCA objects, all of which interface directly with downstream visualization functions.

### Visualization, diagnostics, and sensitivity analysis

SurvMarker provides companion functions for diagnosing PCA structure, visualizing survival-associated signal, and assessing feature stability. Functions such as `plot_scree()` and `plot_cumvar()` summarize variance explained and cumulative variance across PCs, supporting transparent and reproducible PC selection (Figure 1A-B). Survival-associated latent structure can be visualized using `plot_pc12()` and `plot_top2_survival_pcs(),` which display PC scores colored by clinical or molecular annotations (Figure 1C-D).

Feature-level inference is further supported by `plot_null_vs_observed(),` which contrasts observed feature score with its empirical null distribution. Significant features exhibit scores in the extreme tails of the null distribution (Figure 1E), whereas non-significant features show close agreement with the null (Figure 1F). All visualizations support flexible annotation and color mapping, allowing users to tailor graphical outputs while maintaining consistent representation across plots.

To evaluate robustness to PC choice, SurvMarker supports sensitivity analyses via `run_survival_pca_multi_pc(), which` applies the scoring framework across a user-specified grid of PC counts and returns the resulting feature sets. Feature overlap and stability can be visualized using `plot_venn()`(Figure 1G), and `plot_feature_set_tradeoff()` summarizes the relationship between cumulative variance explained and feature set size (Figure 1H). In the example dataset, the number of selected genes decreases as more PCs are included, reflecting a strengthening empirical null model and increasingly stringent calibration of feature scores. Features retained under larger PC sets therefore represent robust contributors to survival-associated latent structure.

### Comparison with fixed-threshold selection

To demonstrate the practical advantages of the proposed framework, we applied SurvMarker to gene expression profiles from the TCGA-LAML cohort (dbGaP accession: phs000178) and evaluated performance using the previously defined 19-gene and 16-miRNA prognostic panels. Unlike the earlier fixed per-PC selection strategy, SurvMarker integrates information across all survival-associated PCs through a unified, statistically calibrated feature score.

Supplementary Figure 2 illustrates that SurvMarker produces a more compact and stable feature sets that is substantially less sensitive to the number of PCs considered. Across Top10, Top20, Top30, and Top50 PC settings, 11 of the 19 prognostic genes and 12 of the 16 prognostic miRNAs are consistently retained, defining a robust core set. Additional genes are incorporated gradually as PC count increases, reflecting progressive refinement rather than abrupt changes driven by analysis choices. In contrast, the fixed-threshold approach yields gene sets whose size increases rapidly with the number of PCs, complicating biological interpretation and limiting reproducibility.

Together, these results demonstrate that SurvMarker improves parsimony, stability, and robustness in feature selection while preserving prognostic signal. By unifying survival analysis, PCA-based feature scoring, empirical null inference, and diagnostic visualization into a single, coherent workflow, SurvMarker provides a practical and reproducible solution for biomarker discovery in high-dimensional survival studies. The package is fully documented with a strong emphasis on transparency, reproducibility, and ease of use. Sensible default settings enable rapid application to new datasets, while flexible user options allow fine-tuning of PC selection, FDR thresholds, covariate adjustment, and visualization preferences. A comprehensive reference manual detailing all functions, arguments, defaults, and plotting options is provided in the Supplementary Material, ensuring that SurvMarker can be readily adapted to diverse high-dimensional datasets and facilitating the robust translation of transcriptomic signals into clinically interpretable prognostic features.

## Acknowledgements

SurvMarker was developed by building on the extensive R ecosystem for statistical modeling and visualization. Core functionality relies on `prcomp` (R Core Team, 2024) for principal component analysis, the `survival` (Therneau, 2024) package for Cox proportional hazards modeling, and widely used visualization tools including `ggplot2` (Wickham, 2016), and `VennDiagram` (Chen et al., 2011) for overlap and stability visualizations. We thank Drs. Timothy J. Ley and Christopher A. Miller (Washington University); Richard Corbett, and Drs. Emilia Lim and Marco Marra (University of British Columbia) for their invaluable assistance in acquiring the gene and miRNA datasets. We thank the broader open-source R community for maintaining the foundational packages that made this work possible.

