## Supplementary Materials for "SurvMarker: An R Package for Identifying Survival-Associated Molecular Features Using PCA-Based Weighted Scores"

### Supplementary Figures

**Figure S1. PCA-based weighted feature scoring framework for survival analysis implemented in SurvMarker.**

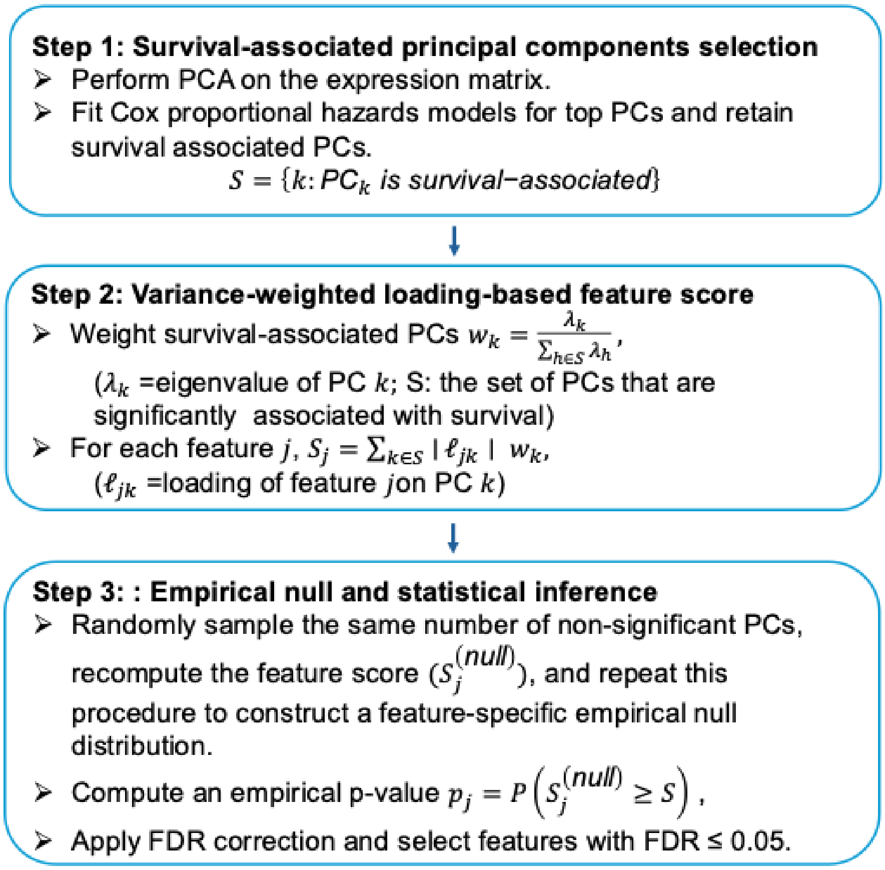

**Figure S2. Comparison of PCA-based weighted feature scoring with fixed per-PC threshold selection.**

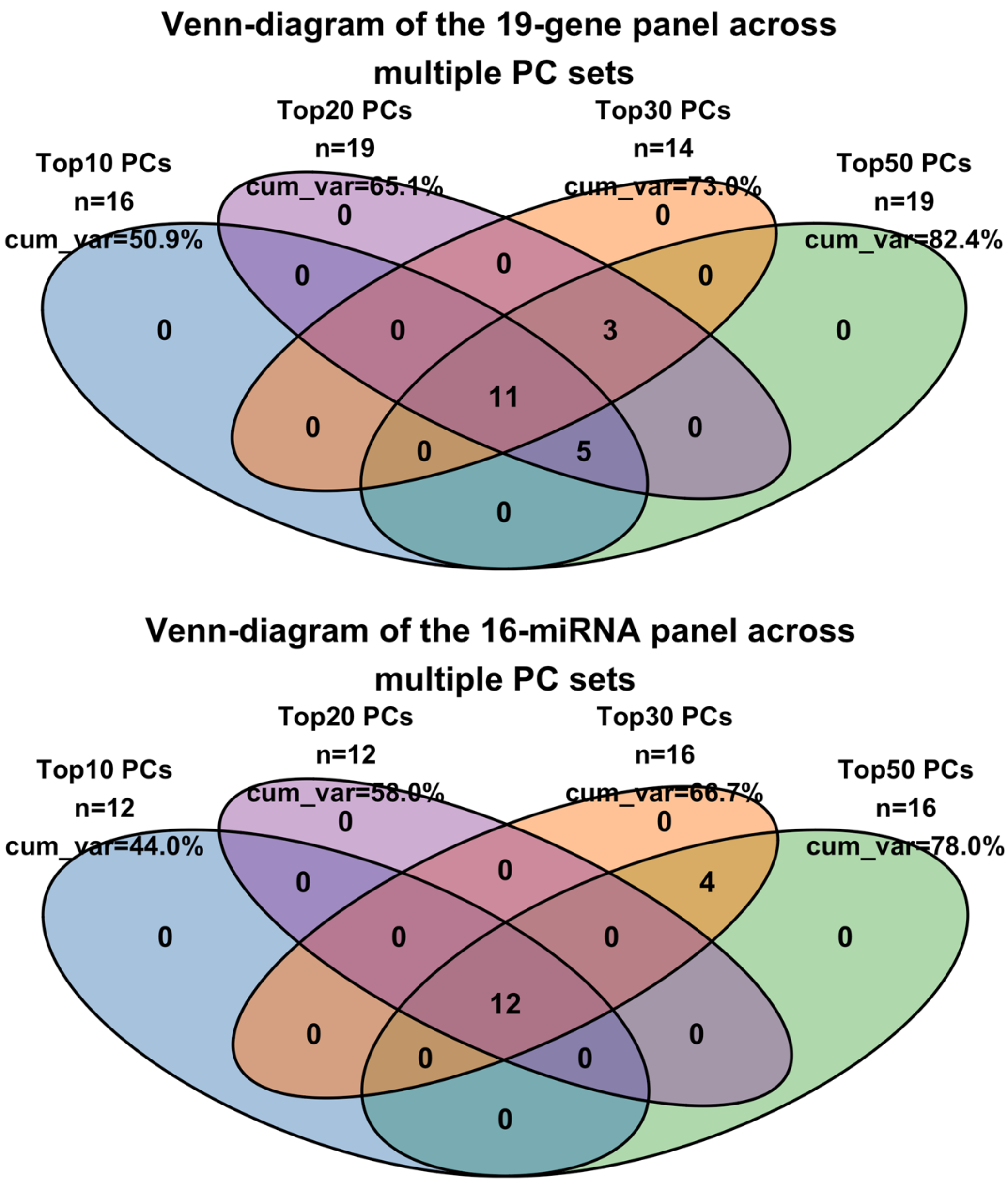

### SurvMarker Reference Manual

SurvMarker: An R Package for Identifying Survival-Associated Molecular Features Using PCA-Based Weighted Scores

#### Description

SurvMarker implements a PCA-based weighted feature-scoring framework for survival analysis that identifies prognostically relevant molecular features by aggregating information across survival-associated principal components and assessing feature significance using an empirical null distribution.

#### Dependencies

`survival`, `ggplot2`, `VennDiagram`

#### 1. PCA-based Feature Scoring

**`pca_based_weighted_score()`**

Purpose: Performs survival-guided PCA to compute weighted feature scores, construct feature-specific empirical null distributions, and identify prognostically significant features. Feature scores aggregate consistent contributions across multiple survival-associated principal components, avoiding arbitrary per-component thresholds. Empirical *p*-values quantify score extremeness relative to null distributions derived from PCs with no survival association.

#### Usage

```
pca_based_weighted_score(
  X,
  time,
  status,
  covar = NULL,
```

```
  n_pcs = 50,
  cumvar_threshold = NULL,
  max_pcs = 50,
  pc_fdr_cutoff = 0.05,
  feature_fdr_cutoff = 0.05,
  null_B = 500,
  seed = 1,
  scale_pca = TRUE,
  use_abs_loadings = TRUE,
  store_null = TRUE,
  verbose = TRUE
)
```

**Arguments**

| | |
|---|---|
| `X` | Numeric matrix or data frame of dimension *n_samples* × *p_features*. Rows correspond to samples and columns to molecular features (e.g., genes, miRNAs, proteins). |
| `time` | Numeric vector of survival times, in the same order as the rows of X. |
| `status` | Numeric or logical event indicator (1 = event, 0 = censored). |
| `covar` | Optional data frame of clinical covariates to include in Cox regression models. |
| `n_pcs` | Integer specifying the number of principal components to retain. Ignored if `cumvar_threshold` is provided. |
| `cumvar_threshold` | Numeric value in (0,1]. Selects the minimum number of PCs required to reach this cumulative variance threshold. |
| `max_pcs` | Hard upper bound on the number of PCs used (safety cap). |
| `pc_fdr_cutoff` | False discovery rate cutoff for selecting survival-associated PCs. |
| `feature_fdr_cutoff` | False discovery rate cutoff for selecting prognostic features. |
| `null_B` | Number of empirical null resamples used for feature-level inference. |
| `seed` | Random seed for reproducibility. |
| `scale_pca` | Logical; whether to scale features prior to PCA. |
| `use_abs_loadings` | Logical; whether to use absolute loadings in feature score calculation. |
| `store_null` | Logical; whether to store the full empirical null score matrix. |
| `verbose` | Logical; whether to print progress messages. |

**Value**

A list containing:

| | |
|---|---|
| `feature_table` | Data frame containing feature loadings on survival-associated PCs, aggregated score ($S_j$), empirical p-values, and false discovery rates (FDR). |
| `pc_table` | PCA summary table including eigenvalues, proportion and cumulative variance explained, Cox regression coefficients, and adjusted p-values for each PC. |

| | |
|---|---|
| `pc_scores` | Sample-level principal component coordinates used for downstream visualization and clustering. |
| `null_scores` | Empirical null score matrix (features × `null_B`) returned when `store_null = TRUE`. |
| `selected_features` | Character vector of prognostically significant features passing feature-level FDR control. |

**Details**

Let $\mathbf{X} \in \mathbb{R}^{n \times p}$ denote the normalized expression matrix with $n$ samples and $p$ features. PCA is applied to $\mathbf{X}$ (optionally after scaling), yielding principal component scores

$$\mathbf{Z} = \mathbf{X}\mathbf{V},$$

where $\mathbf{V} = (v_{jk})$ contains the PC loading vectors and $Z_{ik}$ denotes the score of sample $i$ on PC $k$.

**Identification of survival-associated principal components**

For each retained principal component $k$, its association with survival is evaluated using a Cox proportional hazards model

$$\lambda_i(t \mid Z_{ik}, \mathbf{c}_i) = \lambda_0(t) \exp\left(\beta_k Z_{ik} + \boldsymbol{\gamma}^\top \mathbf{c}_i\right),$$

where $Z_{ik}$ denotes the score of individual $i$on principal component $k$, $\mathbf{c}_i$represents optional clinical covariates (e.g., age, sex, ELN-2022 risk group), $\beta_k$ is the PC-specific log hazard ratio, and $\lambda_0(t)$ is the baseline hazard function. *p*-values are adjusted for multiple testing across PCs using the Benjamini–Hochberg procedure, and PCs with adjusted *p*-values less than `pc_fdr_cutoff` are declared survival-associated.

**Feature score aggregation**

For each feature $j$, a weighted feature score $S_j$ is computed by aggregating its loadings across all survival-associated PCs:

$$S_j = \sum_{k \in \mathcal{K}} w_k \; \ell_{jk},$$

where $\mathcal{K}$ denotes the set of survival-associated PCs, $\ell_{jk} = \mid v_{jk} \mid$ if `use_abs_loadings = TRUE` (or $v_{jk}$ otherwise), and

$$w_k = \frac{\lambda_k}{\sum_{k \in \mathcal{K}} \lambda_k}$$

is a variance-based weight proportional to the eigenvalue $\lambda_k$ (variance explained) of PC $k$. The resulting score $S_j$ reflects the overall contribution of feature $j$ to survival-relevant latent structure.

**Empirical null distribution and inference**

Statistical significance of $S_j$ is evaluated using an empirical null distribution constructed by repeatedly sampling PCs that are not associated with survival. For each null iteration $b = 1, \dots, B$, a null score

$$S_j^{(b)} = \sum_{k \in \mathcal{K}^{(b)}} w_k^{(b)} \ \ell_{jk}^{(b)}$$

is computed using the same aggregation scheme, where $\mathcal{K}^{(b)}$ is a randomly selected set of non–survival-associated PCs of the same size as $\mathcal{K}$.

The empirical p-value is defined as

$$p_j = \frac{\sum_{b=1}^{B} \mathbb{I}\left( S_j^{(b)} \geq S_j \right)}{B},$$

and feature-level false discovery rates are controlled using the Benjamini–Hochberg procedure at level `feature_fdr_cutoff`.

### 2. Visualization Functions

#### 2.1 PCA Variance Diagnostics

**`plot_scree()`**

**Description**

Plots the proportion of variance explained by each PC, with optional annotation of a selected PC cutoff.

**Usage**

```
plot_scree(
  res,
  n_pcs = 50,
  show_threshold = FALSE,
  threshold_pc = NULL,
  ...)
```

**Arguments**

| | |
|---|---|
| `res` | Result object returned by `survival_based_loading_score()`. |
| `n_pcs` | Number of PCs to display. |
| `show_threshold` | Logical; draw a vertical line at `threshold_pc`. |

`threshold_pc` PC index for annotation.

#### Value

A `ggplot` object.

#### `plot_cumvar()`

#### Description

Plots cumulative variance explained across PCs with optional threshold annotation.

#### Usage

```
plot_cumvar(
  res,
  n_pcs = 30,
  show_threshold = FALSE,
  cum_threshold = 0.5,
  ...
)
```

#### Details

Facilitates principled selection of PC dimensionality.

### 2.2 PCA Scatter Plots

#### `plot_pc12()`

#### Description

Creates a scatter plot of two selected principal components with optional coloring and shaping by metadata.

#### Usage

```
plot_pc12(
  res,
  meta = NULL,
  id_col = "PATIENT_ID",
  pc_x = 1,
  pc_y = 2,
  color_by = NULL,
  shape_by = NULL,
  point_size = 3,
  ...
)
```

##### Arguments

| | |
|---|---|
| `res` | Result object returned by `pca_based_weighted_score()`. |
| `meta` | Optional metadata data frame used for sample annotation. |
| `id_col` | Column name in meta identifying sample IDs. |
| `pc_x`, `pc_y` | Integer indices of the principal components plotted on the x- and y-axes. |
| `color_by` | Metadata column used to map points to colors. |
| `shape_by` | Metadata column used to map points to shapes. |
| `point_size` | Numeric value controlling point size. |
| `alpha` | Point transparency level (0–1). |
| `title` | Plot title. |
| `...` | Additional `ggplot2` layers, scales, or theme adjustments. |

##### Value

A `ggplot` object.

##### Typical Use

Visualize disease subtypes, clinical risk groups, or molecular classes in PCA space.

#### `plot_top2_survival_pcs()`

##### Description

Automatically identifies and plots the two PCs with the smallest adjusted survival p-values.

##### Usage

```
plot_top2_survival_pcs(
  res,
  meta = NULL,
  color_by = NULL,
  title = "Top 2 Significant PCs",
  ...
)
```

##### Details

Ensures that displayed PCA structure is directly linked to survival relevance.

### 2.3 Empirical Null Diagnostics

**`plot_null_vs_observed()`**

**Description**

Visualizes the empirical null distribution of a feature's score and overlays the observed score.

**Usage**

```
plot_null_vs_observed(
  res,
  feature,
  bins = 30,
  hist_fill = "lightblue",
  obs_line_color = "red",
  ...
)
```

**Arguments**

| | |
|---|---|
| `feature` | Feature name whose observed score is visualized against the empirical null distribution. |
| `bins` | Number of bins used in the histogram of null scores. |
| `hist_fill` | Fill color for the histogram representing the null distribution. |
| `obs_line_color` | Color of the vertical line indicating the observed feature score. |

**Value**

A `ggplot` object.

**Interpretation**

- Significant features show strong right-tail separation
- Non-significant features overlap with the null distribution

**2.4 Multi-PC Stability Analysis**

**`run_survival_pca_multi_pc()`**

**Description**

Runs the survival-associated PCA-based scoring pipeline across multiple PC cutoffs to assess feature stability.

**Usage**

```
run_survival_pca_multi_pc(
  X,
  time,
  status,
  covar = NULL,
```

```
  pcs_to_run = c (10, 20, 30, 50),
  ...
)
```

#### Value

A list containing feature sets, cumulative variance, and settings across PC choices.

### `plot_venn()`

#### Description

Generates a Venn diagram showing overlap of selected feature sets across PC cutoffs.

#### Usage

```
plot_venn(
  pcset_obj,
  pick,
  label_mode = "both",
  ...
)
```

### `plot_feature_set_tradeoff()`

#### Description

Plots the trade-off between selected feature set size and cumulative variance explained.

#### Usage

```
plot_feature_set_tradeoff(
  pcobj,
  ...
)
```

#### Scientific Value

Demonstrates robustness, stability, and interpretability of the selected feature panel as model complexity increases.
